# Human-level recognition of blast cells in acute myeloid leukemia with convolutional neural networks

**DOI:** 10.1101/564039

**Authors:** Christian Matek, Simone Schwarz, Karsten Spiekermann, Carsten Marr

## Abstract

Reliable recognition of malignant white blood cells is a key step in the diagnosis of hematologic malignancies such as Acute Myeloid Leukemia. Microscopic morphological examination of blood cells is usually performed by trained human examiners, making the process tedious, time-consuming and hard to standardise.

We compile an annotated image dataset of over 18,000 white blood cells, use it to train a convolutional neural network for leukocyte classification, and evaluate the network’s performance. The network classifies the most important cell types with high accuracy. It also allows us to decide two clinically relevant questions with human-level performance, namely (i) if a given cell has blast character, and (ii) if it belongs to the cell types normally present in non-pathological blood smears.

Our approach holds the potential to be used as a classification aid for examining much larger numbers of cells in a smear than can usually be done by a human expert. This will allow clinicians to recognize malignant cell populations with lower prevalence at an earlier stage of the disease.

## Introduction

Microscopic examination and classification of blood cells is an important cornerstone of hematological diagnostics [1, 2, 3]. Specifically, morphological evaluation of leukocytes from peripheral blood or bone marrow samples is one of the initial steps in the diagnosis of hematopoietic malignancies such as Acute Myeloid Leukemia (AML) [4, 5]. In particular, the common French-Americal-British (FAB) classification of AMLs strongy relies on cytomorphology [6]. Having been part of routine workup of hematological diagnosis since the 19th century, cytomorphological examination of leukocytes has so far defied automation and is regularly performed by trained human experts. Therefore, cytomorphological classification is tedious and time-consuming to produce, suffering from considerable intra-and inter-observer variation that is difficult to account for, and hard to deliver in situations where trained experts are lacking. Furthermore, it is difficult to reliably correlate with the result of other, intrinsically more quantitative diagnostic modalities such as immunophenotyping or molecular genetics. Reliable, automated differentiation of cell morphology and recognition of malignant cells is also a key prerequisite to allow screening for hematological neoplasms, potentially enabling their earlier detection and treatment.

As cytomorphological examination is based on evaluating microscopic cell images, it can be formulated as an image classification task. Deep convolutional neural networks (CNNs) have proven very successful in the field of natural image classification [7, 8, 9]. Recently, CNNs have been successfully applied to various medical imaging tasks, including skin cancer recognition [10], evaluation of retinal disorders [11] and the analysis of histological sections [12, 13], for example through mitosis detection [14], region of interest detection and analysis [15] or tissue type segmentation [16]. This motivates us to apply CNNs to cytomorphological classification of blood cells, in particular those relevant in AML.

Previous work on leukocyte classification has mainly been focused on feature extraction from cytological images [17, 18]. In that context, lymphoblastic leukemias, where the cytomorphology is less diverse than in the myeloid case, have received more attention [19, 20]. Providing sufficiently many labelled images for deep learning methods to work has proven challenging in medical image analysis, due to restrictions on access to and expense of expert time for providing ground truth annotations [21, 22]. Therefore, most studies have worked on datasets limited in the number of patients included or individual cytological images classified [23, 24]. Hence, applications of CNNs to white blood cell classification have so far been focused on differentiation of specific subtypes such as erythroid and myeloid precursors [23].

Here, we introduce a database comprising 18,365 individual cell images from 200 individuals, and develop a CNN that is able to classify individual cells from peripheral blood smears and judge for malignancy with high accuracy.

## Materials and Methods

We selected peripheral blood smears from 100 patients diagnosed with different subtypes of AML at the Laboratory of Leukemia Diagnostics at Munich University Hospital between 2014 and 2017, and smears from 100 patients found to exhibit no morphological features of hematological malignancies in the same time frame. The study setup was reviewed by the local ethics committee, and consent was obtained under reference number 17-349.

For all selected blood smear images, we followed the workflow depicted schematically in Fig. 1: An area of interest comprising approximately 20 mm^2^ within the monolayer region of the smear was selected from a low-resolution pre-scan, and scanned at 100-fold optical magnification with oil immersion using an M8 digital microscope / scanner (Precipoint GmbH, Freising, Germany). The resulting digitised data consisted of multiresolution pyramidal images of a size of approximately 1 GB per scanned area of interest. A trained examiner experienced in routine cytomorphological diagnostics at Munich University Hospital differentiated physiological and pathological leukocyte types contained in the microscopic scans into the classification scheme shown in Fig. 2B, which is derived from standard morphological categories and was refined to take into account subcategories relevant for the morphological classification of AML, such as bilobed Promyelocytes, which are typical of the FAB subtype M3v [3]. Annotation was carried out on a single-cell basis, and approximately 100 cells were differentiated in each smear. Subimage patches of size 400 x 400 pixels (corresponding to approximately 29*µ*m x 29*µ*m) around the annotated cells were extracted without further cropping or filtering, including background components such as erythrocytes, platelets or cell fragments. When examining the screened blood smears, the cytologist followed the routine clinical procedure. Overall, 18,365 single-cell images were annotated and cut out of the scan regions. The full class-wise statistics of the dataset are given in Tab. S1, and sample images of the most important physiological and pathological classes are shown in Fig. 2A and C. The database of single-cell images is provided online.

**Figure 1:**
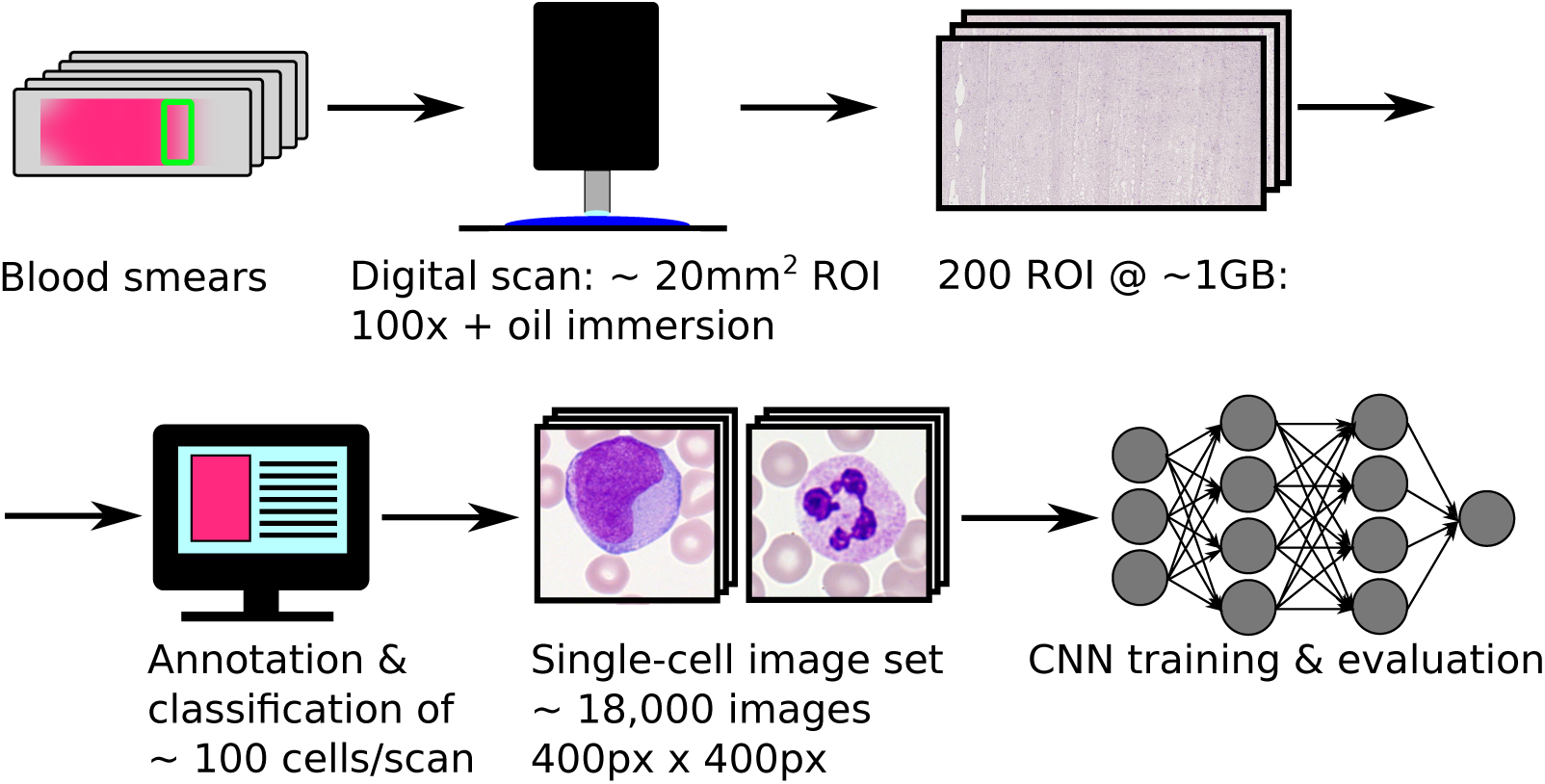
Data handling workflow. Peripheral blood smears from 100 AML patients and 100 patients without signs of hematological malignancy were digitised using an oil-immersion microscope at 100-fold magnification. After annotation of single cells using the classification scheme shown in Fig. 2B, a convolutional neural network was trained and evaluated.

**Figure 2:**
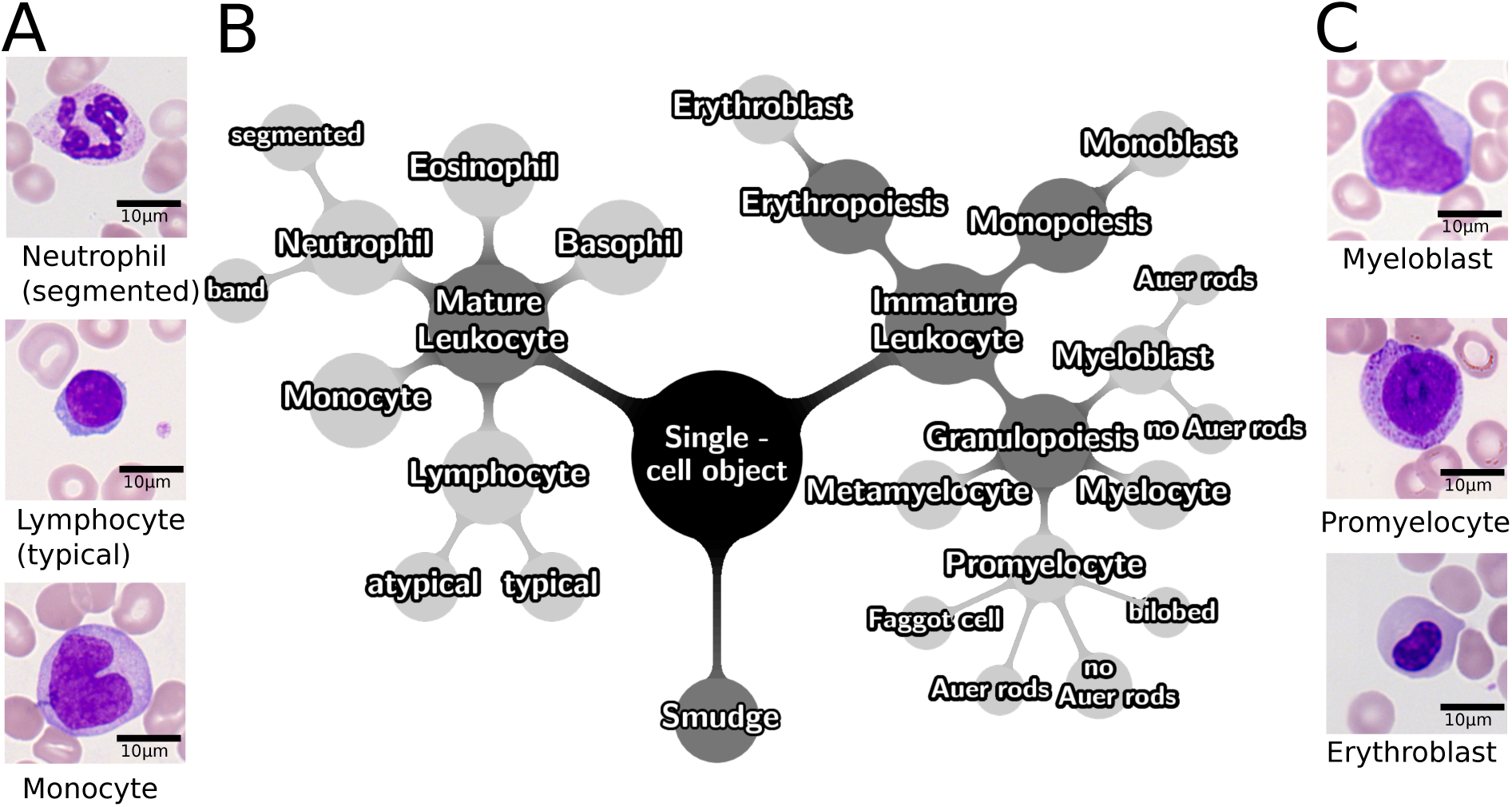
Classification of 18,000 single-cell images into a 18-class scheme. (A): Sample images of the three most frequent mature cell classes contained in the dataset. Scale bars correspond to 10 *µm*. (B): Taxonomy tree of the classification scheme used for annotation of single-cell images. Cell morphologies present under physiological conditions are contained in the left branch, while cells normally absent under physiological conditions are contained in the right branch. Smudge cells are classified in a separate category. To ensure sufficient population of classes, some leaves of the taxonomy were combined into overall classes for training and testing (cf. main text). (C): Sample images of the three most frequent immature cell classes contained in the dataset. Scale bars correspond to 10 *µm*.

Annotations of single-cell images provide the ground truth for training and evaluation of our network. Morphological classes containing fewer than 10 images were merged with neighbouring classes of the taxonomy. Specifically, myeloblasts with and without Auer rods were merged into a common myeloblast class, and faggot cells and promyelocytes with and without Auer rods were merged into a common promyelocyte class, resulting in 15 classes for training and evaluation. A subset of 1,905 single-cell images from all morphological categories were presented to a second, independent examiner, and annotated for a second time in order to estimate inter-rater variability.

For our image classification task, we used a ResNeXt CNN topology described by Xie *et al.* [25], which derives from a residual network, and achieved a second rank in the classification task of the ImageNet ILSVRC 2016 competition. While several versions of residual networks have been shown to be successful in natural image classification, ResNeXt is characterised by a comparatively small number of free hyper-parameters, and is therefore expected to be a convenient choice, in particular as no networks pre-trained on similar datasets are available. We adopted the network to input image dimensions of 400 x 400 x 3 and retained the cardinality hyper-parameter at *C* = 32 as used in Ref. [25], avoiding further tuning of hyper-parameters based on our dataset. The ResNeXt network was implemented using the implementation of Ref. [26] for Keras 2.0 [27], with the input size adjusted to accept images of size 400*x*400 pixels, and the final dense layer adapted to our 16-category classification scheme.

The network was trained for at least 20 epochs, which took a computing time of approximately 4 days on a Nvidia GeForce GTX TITAN X GPU.

We randomly divide the images contained in each class of our dataset in a test-and training group, where the training group contains approximately 80%, and the test group 20% of the images. For 5-fold cross-validation, we performed a stratified split of the cell images into 5 folds, where each fold contains approximately 20% of the images in each class. An individual image is contained in one fold only. Consequently, five different models are trained, each of which uses one fold for testing, and the four remaining folds for training.

In order to cope with the imbalance of cell numbers contained in different classes and take advantage of the rotational invariance of the cell classification problem, we generated additional images by applying random rotational transformations of 0 – 359 degrees, as well as random horizontal and vertical flips to the single-cell images in our dataset. Using these operations, we augmented the data set in such a way that each class contained approximately 10,000 images for training.

To quantitatively evaluate the class-wise accuracy, we calculate precision, specificity and sensitivity as comparison metrics, which are defined as follows:

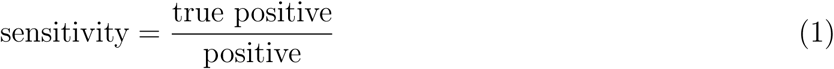

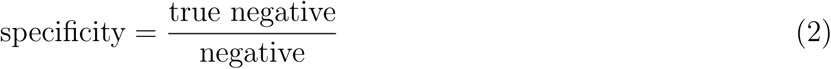

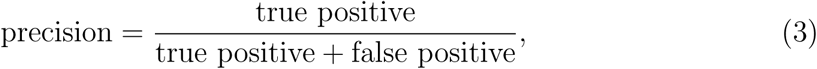

where “true positive” and “true negative” are the number of images correctly ascribed or not ascribed to a given class by the network, respectively, “positive” and “negative” are the overall number of images shown belonging or not belonging to a certain class, and “false positive” is the number of cell images wrongly ascribed to that class.

## Data and code availability

Code for the network trained in this study and network weights are avaiable at CodeOcean, together with a subset of the single-cell image data used to test the network. Additional image data may be available from the authors upon reasonable request, with permission from Munich University Hospital.

## Results

We evaluate the performance of the trained network by feeding single-cell images through it, and comparing the output prediction with the labels assigned by human examiners. The network outputs a vector of probabilities **P** = (*P*_1_, *…, P*_*i*_, *…, P*_15_), where the components *P*_*i*_ are the respective predicted probabilities for the image to belong to class *i* out of the 15 overall classes. The network’s image class prediction is then the class *m* with the highest corresponding predicted probability *P*_*m*_.

The class-wise prediction accuracy of the network is shown in the confusion matrix of Fig. 3A. We note that the network achieves excellent agreement with human annotations (each above 90%) for the most common physiological cell types, including segmented neutrophils, typical lymphocytes, monocytes, and eosinophils. Also myeloblasts, whose presence in the peripheral blood is common in myeloid leukemias [5], are recognised with high accuracy, yielding a precision and sensitivity of 94% (cf. Tab. 1). Other classes are more challenging for the network, in particular the intermediate stages of granulopoiesis and erythropoiesis, and basophils,where our test and training dataset contains less than 100 images. Values of precision and sensitivity for all cell classes obtained by 5-fold cross-vaildation are given in Tab. 1. We note that due to the varying number of specific cell types present in smears, the number of test and training images varies by up to two orders of magnitude for different classes. In order to avoid biased evaluation of our classifier, we refer to class-wise precision and sensitivity, and do not evaluate an overall accuracy score, which would be biased towards the classes with a high number of samples [7].

**Table 1:**
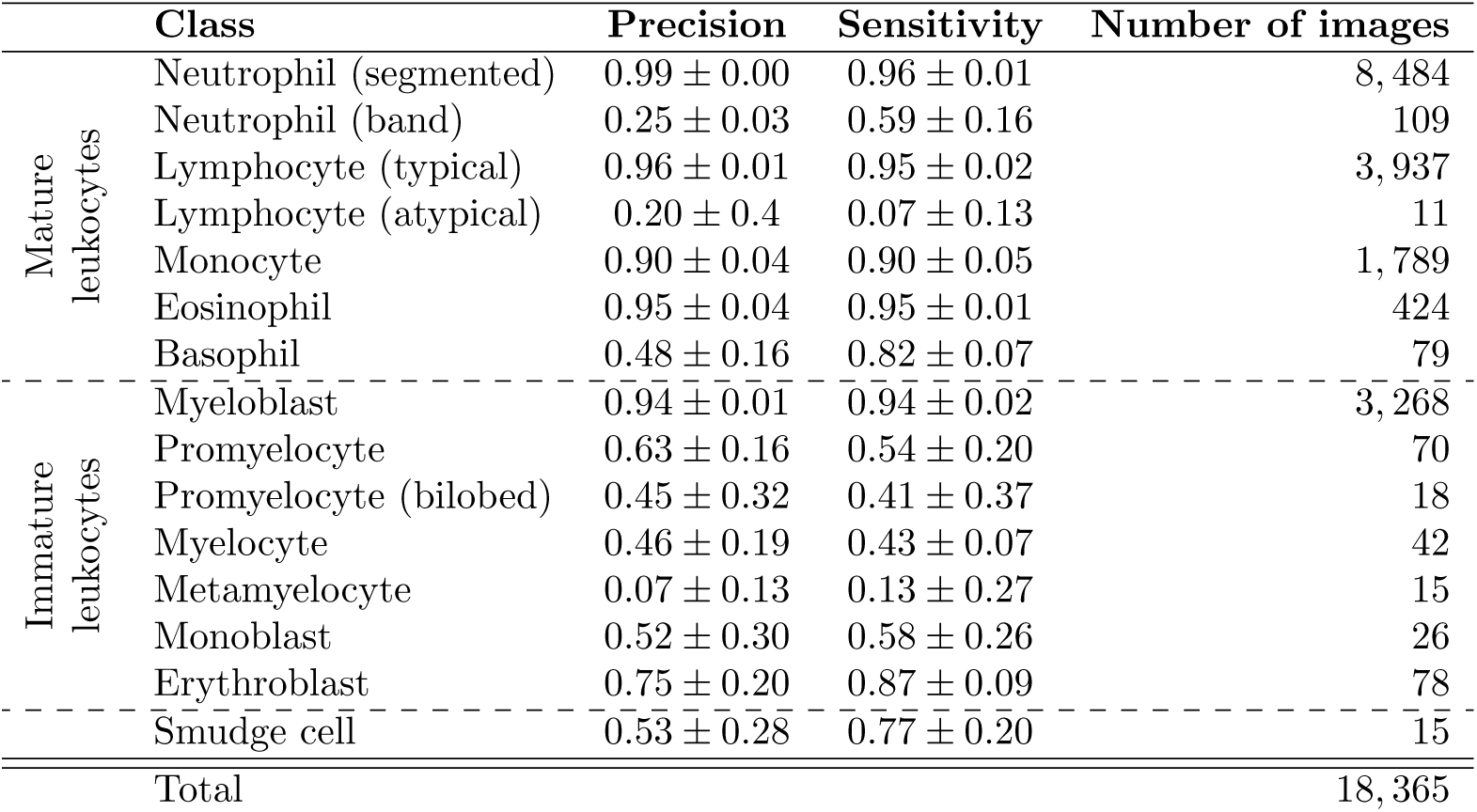
Class-wise precision and sensitivity of the network, determined by 5-fold cross-validation. The model achieves precision and sensitivity above 0.9 on classes for which more than 100 images are available, such as segmented neutrophils, typical lymphocytes and myeloblasts. Large deviations across folds occur for classes with small sample number, e.g. metamyelocytes and promyelocytes.

**Figure 3:**
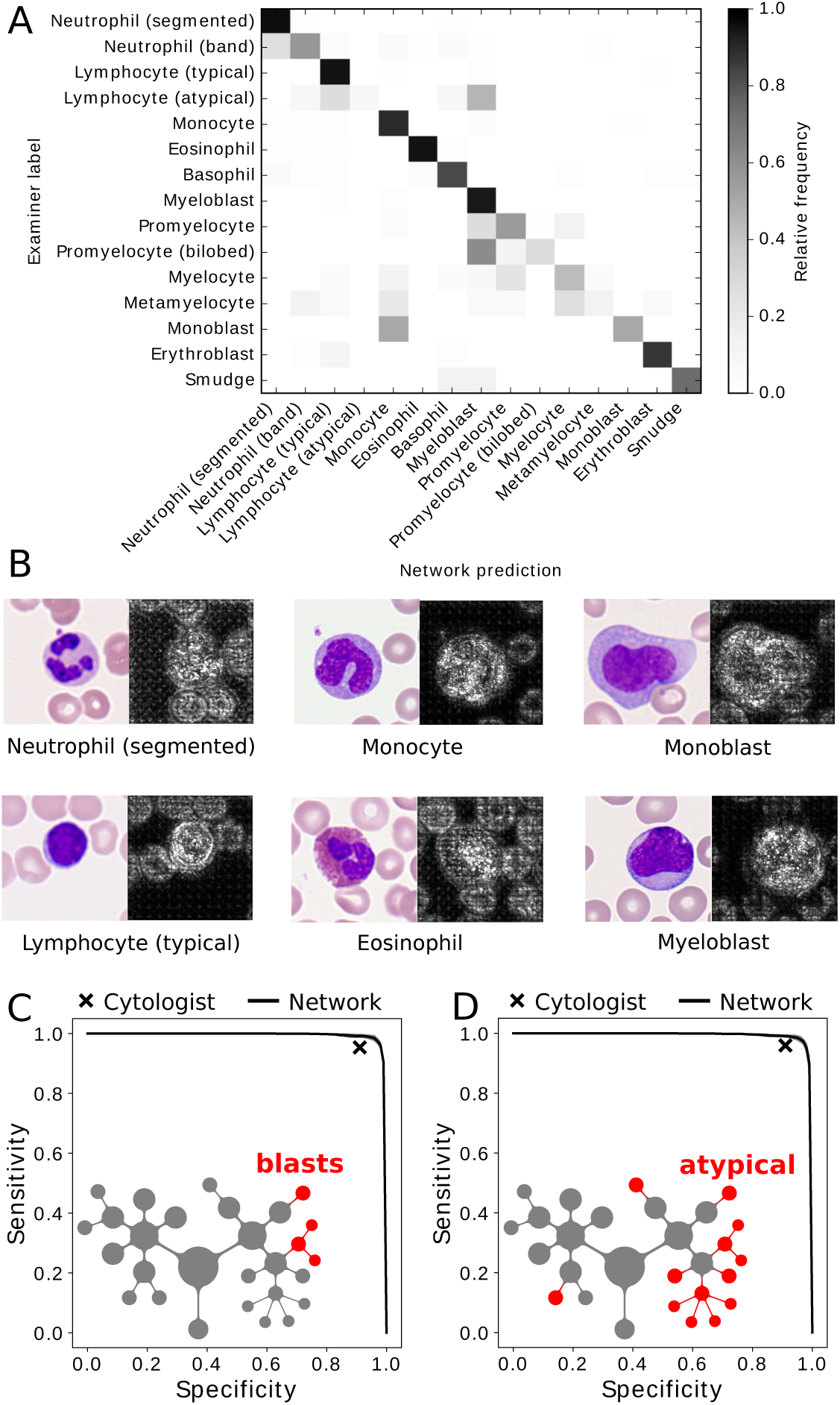
Human-level network performance in single-cell classification, pixel-wise attention and binary decision tasks. (A): Class-wise agreement between network prediction and ground-truth human examiner label. Outstanding classification performance is observed for key cell classes such as myeloblasts. The full, class-wise statistics of prediction quality is given in Tab. 1. (B): Saliency maps illustrate the gradient of a pixel with respect to the networks’s loss function. Brighter pixels have a higher influence on the network’s classification decision. Maps suggest that the network has learned to focus on the leukocyte and map out its internal structures, while giving less weight to background content. (C): ROC curve of binary decision between cells with and without blast character. The network performs very well with an AUC of 0.992 *±* 0.001 (obtained by training *n* = 5 networks for cross-validation, each tested on a separate 20% of overall image data), and attains the performance of a second human annotator shown by ‘x’. Inset: Schematic depiction of cell class taxonomy, where blast equivalent classes are coloured red. (D): On the binary classification for atypical cells normally absent in non-pathological smears, outstanding performance is observed with an AUC of 0.991 *±* 0.002. Also for this binary question, the network attains human-level performance. Inset: Schematic depiction of cell class taxonomy, with atypical cell classes coloured red.

In order to relate the network performance to the inter-rater variability encountered among human examiners, we asked another, independent cytologist to re-annotate a subset of 1,905 single-cell images containing all subtypes previously annotated. The level of agreement between the two annotators is shown in Fig. S1. Notably, the CNN and the second annotator show similar patterns of deviation to the first examiner, specifically as far as classification of atypical lymphocytes and promyelocytes as myeloblasts is concerned. This may reflect visual similarities between instances of these cell types recognised by both the network and human examiners, which intrinsically limit the single-cell labelling process. Furthermore, the first examiner had access to the whole smear and was hence able to compare to other cells present, whereas the second examiner, just as the network, classified only single-cell images.

To evaluate if our network fixates on relevant parts of the single-cell images, we calculated saliency maps following the procedure outlined by Simonyan *et al.* [28]. These maps allow visualising how important a given pixel is for the network’s classification decision. Saliency maps for several test images are shown in Fig. 3B, demonstrating that pixels within the leukocyte are most important for the network’s classification decision, suggesting that the network has learned to focus on relevant areas of the input image.

A key clinical question when examining blood cell morphology is if a given cell is a myeloblast or monoblast, as these two cell types are counted as blast equivalents, and are generally required to be present in the peripheral blood for a diagnosis of AML [5]. Using the output of our network, we can determine the probability of a cell to possess blast character, *P*_blast_ = *P*_myeloblast_ + *P*_monoblast_, and choose a threshold probability *t*, so that the binary prediction of the network is given by *ŷ* = *P*_blast_ ≥*t*. The receiver operating characteristic (ROC) curve is the result of sweeping t between 0 and 1, and is shown in Fig. 3C. The area under the curve (AUC), which we measure as 0.992 ± 0.001 using 5-fold cross-validation, shows that out network provides a test of the blast character of a given single-cell image that can be considered outstanding by the criteria of diagnostic test assessment [29]. In comparison to the network’s performance, the second human rater reproduces the cell label provided by the first annotator with a sensitivity of 95.3% and a specificity of 91.1% (cf. Fig. 3C), which lies close but somewhat below the network ROC curve.

Another clinically important, binary decision on individual white blood cells is whether a given cell belongs to one of the typical cell types, present in peripheral blood under normal circumstances, or to atypical cell types that occur in pathological situations, namely myeloblasts, monoblasts, myelocytes, metamyelocytes, promyelocytes, erythroblasts and atypical lymphocytes. As in the test for blast character, we determine the overall probability *P*_atypical_ for a given cell to belong to one of these groups by adding the output probabilities of all atypical cell classes, and defining a threshold probability *t* for the atypicality test to be positive. Again, the network yields an outstanding test of the atypicality of a cell in a given image, with an AUC of 0.991 ± 0.002, which compares to a human second-rater sensitivity of 95.9% and specificity of 91.0% (Fig. 3D). Importantly, our network outperforms the cytologist who re-annotated the subset of 1,905 single-cell images in both clinically relevant binary decision tasks tested here (cf. Fig. 3C and D).

## Discussion

The convolutional neural network presented in this study shows Outstanding performance at identifying the most important morphological white blood cell types present in non-pathological blood, as well as the key pathological cell types in Acute Myeloid Leukemias. For the most common physiological leukocyte classes as well as for myeloblasts, it attains precision and sensitivity values above 90%, allowing these cells to be identified with a very high accuracy which outperforms other classifiers in the literature [23, 18]. The classification predictions can be used to answer clinically relevant binary questions. Blast character and atypicality are of high relevance in practice and can be provided with very high confidence by the network.

As expected for our data-driven classification method, a correlation can be observed between the number of images available for a specific class in our data set and the performance of the network on that class (cf. Tab. 1). Although to the authors’ knowledge, the image set presented in this paper is the largest used so far in the literature, we anticipate that further enlarging the dataset would improve the network’s classification performance also for rare cell types.

Sources of disagreement between the network and the ground truth are linked to the inter-rater variability of the cytomorphologic examination, which is known to limit the reproducibility particularly of rare leukocyte species [30]. We have estimated inter-observer variability of cytomorphologic classification in our dataset by re-annotation of single-cell images. We note that the inter-rater variability is particularly high for myeloblasts, reflecting to some degree the polymorphic nature of that cell class. Furthermore, the examiner providing the ground-truth labels for single-cell images had access to the whole scan and was therefore able to compare the morphologies of cells present in a smear. In contrast, both the second annotator and the network had to make a classification decision based solely on a single-cell image, without the ability to compare to other cells from the same patient.

We have compiled a dataset of 18,365 single-cell images of different cell morphologies relevant in the diagnosis of AML from the peripheral blood smears of 200 individuals. After annotation by human experts, we used this dataset to train and evaluate a state-of-the-art image classification convolutional neural network. The network shows good performance at differentiating morphological cell types important for recognising malignancy in peripheral blood smears. For the diagnostically relevant binary questions if a given cell is considered to have blast character or to be atypical, the network achieves outstanding accuracies with an ROC AUC of approximately 0.99 in both cases. Given this level of performance and the fact that our method is scalable and fast, our algorithm can be used to quickly evaluate thousands of cells on a blood smear scan, helping cytologists to find suspicious cells more readily. This might be particularly useful in situations where the number of malignant cells is expected to be small, such as in the early stages of the disease or beginning relapse. In our present paper, we tested the model on peripheral blood smears from one lab scanned with one type of scanning device. In order to evaluate the model’s performance in a realistic routine setting in more detail, further validation is required using data from different sources and disease classes. However, given the variability already present in a dataset compiled from 200 independently stained smears from different patients, we expect the variability to be represented reasonably well in our dataset.

Our method holds the potential to act as a rapid pre-screening and quantitatively informed decision tool for cytological examiners, and might further increase its performance when combined with additional, intrinsically quantitative methods used in the diagnosis of hematological malignancies, such as flow cytometry or molecular genetics.

## Acknowledgements

We thank Nikos Chlis for comments on the manuscript. This work was supported by German Research Foundation grant SFB 1243. Christian Matek gratefully acknowledges support from Deutsche José Carreras-Leukämie Stiftung.

